# Comprehensive pathway analyses of schizophrenia risk loci point to dysfunctional postsynaptic signaling

**DOI:** 10.1101/117481

**Authors:** Dick Schijven, Daniel Kofink, Vinicius Tragante, Marloes Verkerke, Sara L. Pulit, René S. Kahn, Jan H. Veldink, Christiaan H. Vinkers, Marco P. Boks, Jurjen J. Luykx

## Abstract

Large-scale genome-wide association studies (GWAS) have implicated many low-penetrance loci in schizophrenia. However, its pathological mechanisms are poorly understood, which in turn hampers the development of novel pharmacological treatments. Pathway and gene set analyses carry the potential to generate hypotheses about disease mechanisms and have provided biological context to genome-wide data of schizophrenia. We aimed to examine which biological processes are likely candidates to underlie schizophrenia by integrating novel and powerful pathway analysis tools using data from the largest Psychiatric Genomics Consortium schizophrenia GWAS (N = 79 845) and the most recent 2018 schizophrenia GWAS (N = 105 318). By applying a primary unbiased analysis (Multi-marker Analysis of GenoMic Annotation; MAGMA) to weigh the role of biological processes from the MSigDB database, we identified enrichment of common variants in synaptic plasticity and neuron differentiation gene sets. We supported these findings using MAGMA, Meta-Analysis Gene-set Enrichment of variaNT Associations (MAGENTA) and Interval Enrichment Analysis (INRICH) on detailed synaptic signaling pathways from the Kyoto Encyclopedia of Genes and Genomes (KEGG) and found enrichment in mainly the dopaminergic and cholinergic synapses. Moreover, shared genes involved in these neurotransmitter systems had a large contribution to the observed enrichment, protein products of top genes in these pathways showed more direct and indirect interactions than expected by chance, and expression profiles of these genes were largely similar among brain tissues. In conclusion, we provide strong and consistent genetics and protein-interaction informed evidence for the role of postsynaptic signaling processes in schizophrenia, opening avenues for future translational and psychopharmacological studies.

## 1. Introduction

Although post-mortem studies, imaging and human genetic studies have contributed to theories about pathophysiological mechanisms in schizophrenia, the underlying molecular processes have not been fully elucidated. This knowledge gap hampers the development of novel pharmacological treatments. Genetic studies provide a valuable resource to investigate the mechanisms that are likely at play in schizophrenia. Schizophrenia is highly heritable (h^2^ estimates ranging from 45-80%) and polygenic (Lichtenstein et al., 2006; Wang et al., 2017). The two largest genome-wide association studies (GWAS) have identified 108 (Schizophrenia Working Group of the Psychiatric Genomics Consortium, 2014) and 145 independent associated risk loci (Pardiñas et al., 2018).

Pathway and gene set enrichment analysis methods are widely used to provide biological context to the results of genetic association studies by testing whether biologically relevant pathways or sets of genes are enriched for genetic variants associated with a phenotype (de Leeuw et al., 2016). These analyses have been widely applied to schizophrenia, providing evidence for the involvement of synaptic and immune-related processes (Duncan et al., 2014; Lips et al., 2012) and insight into possible new drug targets (Gaspar and Breen, 2017). Such findings are supported by pathway analyses in combined psychiatric disorders (schizophrenia, depression and bipolar disorder), revealing enrichment of genetic variants in neuronal, immune and histone pathways (Network and Pathway Analysis Subgroup of Psychiatric Genomics Consortium, 2015). Involvement of calcium signaling and ion channels in schizophrenia has been reported in a gene set analysis paper combining GWAS data with post-mortem brain gene expression data (Hertzberg et al., 2015). Importantly, none of the abovementioned pathway analysis studies has used the full dataset reported in the latest Psychiatric Genomics Consortium schizophrenia GWAS (Schizophrenia Working Group of the Psychiatric Genomics Consortium, 2014). Moreover, several novel or widely used pathway analysis tools have not yet been applied to this schizophrenia GWAS. These tools constitute fast and powerful approaches to test gene set enrichment, despite variability in their correcting for confounding factors that may increase the type 1 error rate (de Leeuw et al., 2016). Additionally, tools and databases aimed at the integration of GWAS data with gene expression and protein-protein interaction data allow to further explore the biological impact of common variants associated with schizophrenia (Lonsdale et al., 2013; Rossin et al., 2011).

Aiming to comprehensively investigate the possible biological processes underlying schizophrenia, we set out to apply gene set and pathway enrichment analysis methods to the 2014 schizophrenia GWAS (Schizophrenia Working Group of the Psychiatric Genomics Consortium, 2014) and additionally integrate the results of these analyses with data on protein-protein interactions and tissue-specific gene expression (Figure 1). We then validated our main findings using the most recent 2018 schizophrenia GWAS (Pardiñas et al., 2018). We thus elucidate the involvement of neuron differentiation and synaptic plasticity in schizophrenia and reveal an accumulation of variants in post-synaptic signaling cascades. They moreover enable a more nuanced understanding of the several actionable classes of neurotransmitters implicated in the disease.

**Figure 1.**
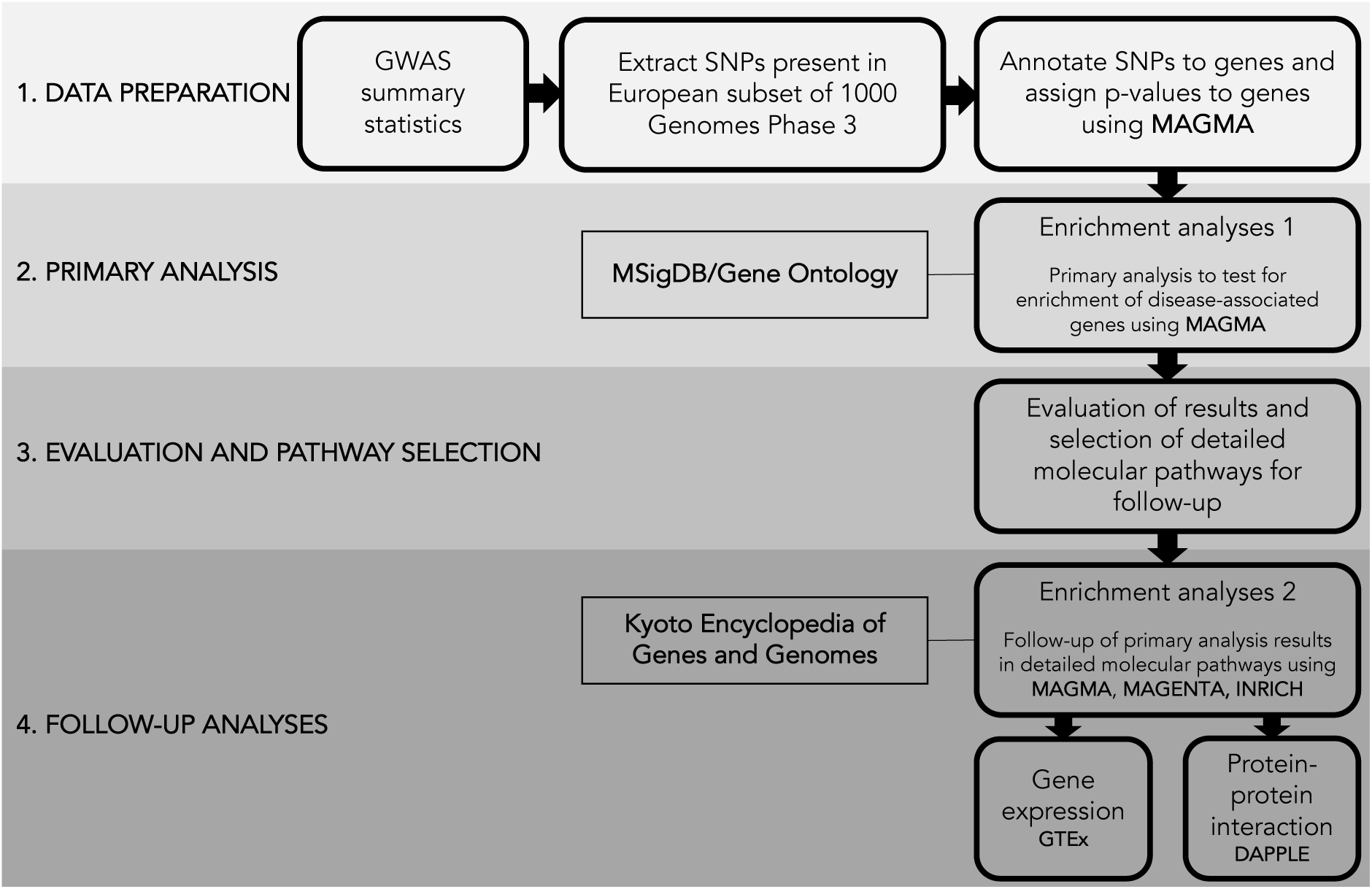
Overview of pathway analysis pipeline. Our analysis pipeline consisted of four stages: **(1)** Data preparation; **(2)** Primary gene set enrichment analysis on MSigDB gene ontology (GO) biological processes; **(3)** Evaluation of primary results and selection of detailed pathways from the Kyoto Encyclopedia of Genes and Genomes (KEGG); **(4)** Follow-up analyses on these detailed molecular pathways to further investigate involvement of biological processes found in the primary analysis.

## 2. Materials and methods

### 2.1 Input data and analysis overview

We used summary-level results from the largest and publically available Psychiatric Genomics Consortium GWAS in schizophrenia (www.med.unc.edu/pgc/results-and-downloads; downloaded on 10 May, 2017) (Schizophrenia Working Group of the Psychiatric Genomics Consortium, 2014). This GWAS was performed in 34 241 schizophrenia cases and 45 604 healthy controls and association results for 9.4 million single-nucleotide polymorphisms (SNPs) were reported in the summary-level data. As detailed below (also see Figure 1), using Multi-marker Analysis of GenoMic Annotation (MAGMA) (de Leeuw et al., 2015) we successively (A) mapped SNPs to genes, (B) calculated gene p-values based on GWAS SNP p-values, (C) performed a primary gene set enrichment analysis using gene ontology (GO) terms, and (D) tested the robustness of these findings in detailed molecular pathways derived from KEGG. We then validated these findings using summary statistics derived from the latest 2018 schizophrenia GWAS (Pardiñas et al., 2018) (downloaded on 1 March, 2018). Finally, we further investigated the results of the analysis on KEGG pathways using Meta-Analysis Gene-set Enrichment of variaNT Associations (MAGENTA) and Interval Enrichment Analysis (INRICH) (Lee et al., 2012; Segrè et al., 2010), applied *in silico* protein-protein interaction (PPI) analysis using Disease Association Protein-Protein Link Evaluator (DAPPLE) (Rossin et al., 2011), and assessed tissue-specific expression using data from the Genotype-Tissue Expression (GTEx) project (Lonsdale et al., 2013).

### 2.2 Mapping SNPs to genes and assigning p-values to genes

SNPs present in the European subset of the 1000 Genomes Phase 3 dataset were extracted from the GWAS summary-level results (The 1000 Genomes Project Consortium, 2015). Using MAGMA v1.06, we mapped SNPs to corresponding genes, extending gene footprints by an additional 20 kilobase (kb) up- and downstream, as a large proportion of regulatory elements involved in gene expression regulation is likely to be captured by including this region (Veyrieras et al., 2008). We then applied a gene analysis to obtain a p-value for each gene to which at least one SNP was mapped. The p-value of a gene was based on the mean association statistic of SNPs contained in that gene. Linkage disequilibrium (LD) between SNPs included in the gene analysis was estimated from the European subset of 1000 Genomes Phase 3, which corresponds best with the ancestry of samples in the schizophrenia GWAS (Schizophrenia Working Group of the Psychiatric Genomics Consortium, 2014).

### 2.3 Gene Ontology (GO) gene set enrichment analysis

We downloaded 4436 GO biological process gene sets from the Molecular Signatures Database (MSigDB, release 6.0, April 2017) (Gene Ontology Consortium, 2015; Liberzon et al., 2011). MSigDB merged original GO terms with high similarity (Jaccard’s coefficient > 0.85) into one gene set and the number of genes per gene set was limited to 10 < n_genes_ < 2000. We applied a competitive gene set analysis, which tests whether the genes in each gene set are more strongly associated with the phenotype of interest than genes outside the gene set, using gene p-values from the mapping step. The significance threshold (at α = 0.05) was adjusted for multiple GO gene sets tested using a permutation procedure implemented in MAGMA (10 000 permutations, p = 4.23×10^-6^). A correction for confounding factors (gene size, gene density and minor allele count) was applied. We additionally tested the robustness of our analysis by implementing several sensitivity analyses: 1) excluding the highly associated MHC region on chromosome 6 (chr6: 25 500 000 - 33 500 000, human genome assembly GRCh37/hg19), 2) excluding the X-chromosome from the analysis, and 3) applying strict filtering for gene set size (10 ≤ n_genes_ ≤ 200) as larger gene sets are sometimes perceived as too broad or uninterpretable.

### 2.4 Follow-up using KEGG pathways

To further test the results of our primary GO gene set analysis, we used pathways representing synaptic signaling processes from the Kyoto Encyclopedia of Genes and Genomes (KEGG, www.kegg.jp, downloaded on 3 January, 2017) (Kanehisa and Goto, 2000) as input for MAGMA: Glutamatergic synapse (hsa04724, 114 genes), cholinergic synapse (hsa04725, 111 genes), serotonergic synapse (hsa04726, 113 genes), GABAergic synapse (hsa04727, 88 genes), dopaminergic synapse (hsa04728, 130 genes), long-term potentiation (hsa04720, 67 genes), and long-term depression (hsa04730, 60 genes). We then ran a similar competitive gene set analysis on these pathways (multiple-testing corrected p = 8.6×10^-3^). For each significantly associated pathway, we mapped top associated genes (gene p < 1.45×10^-4^: 0.05/344 unique genes included in the enrichment analysis of the selected KEGG pathways) to pathway components using *pathview* in R version 3.3.3 (www.r-project.org) (Luo and Brouwer, 2013). Because many genes overlapped between two or more of the tested KEGG pathways (Supplementary Figure S1), we conditioned the enrichment specific to each KEGG pathway on the association signal of genes shared with other tested KEGG pathways (built-in function in MAGMA). Additionally, we tested enrichment of sets of unique genes per KEGG pathway and a gene set containing all shared genes.

To support the results of the MAGMA analysis, we tested enrichment in the same KEGG pathways using MAGENTA and INRICH. MAGENTA calculated gene p-values based on the lowest SNP p-value in a gene (extended 20 kb up- and downstream) while correcting for confounding factors, such as gene length and gene overlap. It then assessed whether a gene set was enriched with low gene p-values at the 95^th^ percentile cut-off (based on all gene p-values in the genome) compared to randomly sampled gene sets of similar size (Segrè et al., 2010). Enrichment p-values were corrected for multiple testing using false discovery rate (FDR < 0.05). INRICH tested whether intervals around associated variants overlap more often with genes in a gene set than could be expected by chance (Lee et al., 2012). Intervals were calculated around genome-wide significant SNPs (p < 5×10^-8^) using the clump function in PLINK v1.90b3z (Chang et al., 2015). Only SNPs with a LD R^2^ > 0.5 and an association p-value < 0.05 were included in an interval, resulting in 114 non-overlapping intervals. Target gene regions were extended with 20 kb up- and downstream. Enrichment of KEGG pathways was tested using a permutation procedure where the enrichment of a gene set was tested against the normal distribution of enrichment in background gene sets. Enrichment p-values were corrected for multiple testing by a bootstrap method in INRICH.

### 2.5. Protein-protein interaction analysis

To increase insight into the biological impact of the pathways implicated in schizophrenia by our analyses, we performed *in silico* PPI analysis using DAPPLE v0.17 (Rossin et al., 2011). We used DAPPLE to test whether proteins in a network comprised of 22 unique top genes from the KEGG dopaminergic, cholinergic or long-term potentiation pathways (gene p < 1.45×10^-4^) show more direct and indirect interactions with each other and with other proteins than expected by chance.

### 2.6 Gene expression analysis

Median relative transcript abundances (transcripts per million, TPM) among 13 brain tissues and 40 non-brain tissues were obtained from the GTEx database (v7, RNASeQCv1.8.8, downloaded on 01-11-2017 from https://www.gtexportal.org/home/datasets) (Lonsdale et al., 2013). Per enriched KEGG pathway, we extracted TPM levels of top genes (gene p < 1.45×10^-4^) and visualized the median relative abundance of transcripts in heat maps for each enriched pathway and for one set of genes that were shared between two or more of the enriched KEGG pathways using the R-package *gplots* (Warnes et al., n.d.). Genes with high similarities in transcript abundance, as well as tissues with similar transcript abundance, were clustered. Additionally, we used median gene expression level in brain (all brain tissues) as a covariate in the MAGMA competitive gene set analysis of two significant GO gene sets and the tested KEGG pathways to assess whether enrichment was dependent on brain gene expression.

## 3. Results

Primary gene set enrichment analysis in MAGMA (in 4436 MSigDB GO biological processes) identified two gene sets that were enriched for schizophrenia-associated SNPs (Figure 2A and Supplementary Table S1): ‘Regulation of Synaptic Plasticity’ (p = 6.52×10^-7^) and ‘Regulation of Neuron Differentiation’ (p = 1.00×10^-6^), which are specialized GO terms of synaptic signaling and neuron differentiation (Supplementary Figure S2). Excluding the extended MHC region or the X-chromosome did not affect the outcome of this analysis. When we applied strict filtering for gene set size, we found that the ‘Regulation of neuron differentiation’ gene set (containing 554 genes) was no longer significant and only ‘Regulation of Synaptic Plasticity’ remained significant (Supplementary Figure S3). There was extensive coherence between the significant GO gene sets (54 shared genes, Supplementary Figure S1B), confirmed by an attenuation of enrichment when the analysis was conditioned on these shared genes (Supplementary Figure S4A).

**Figure 2.**
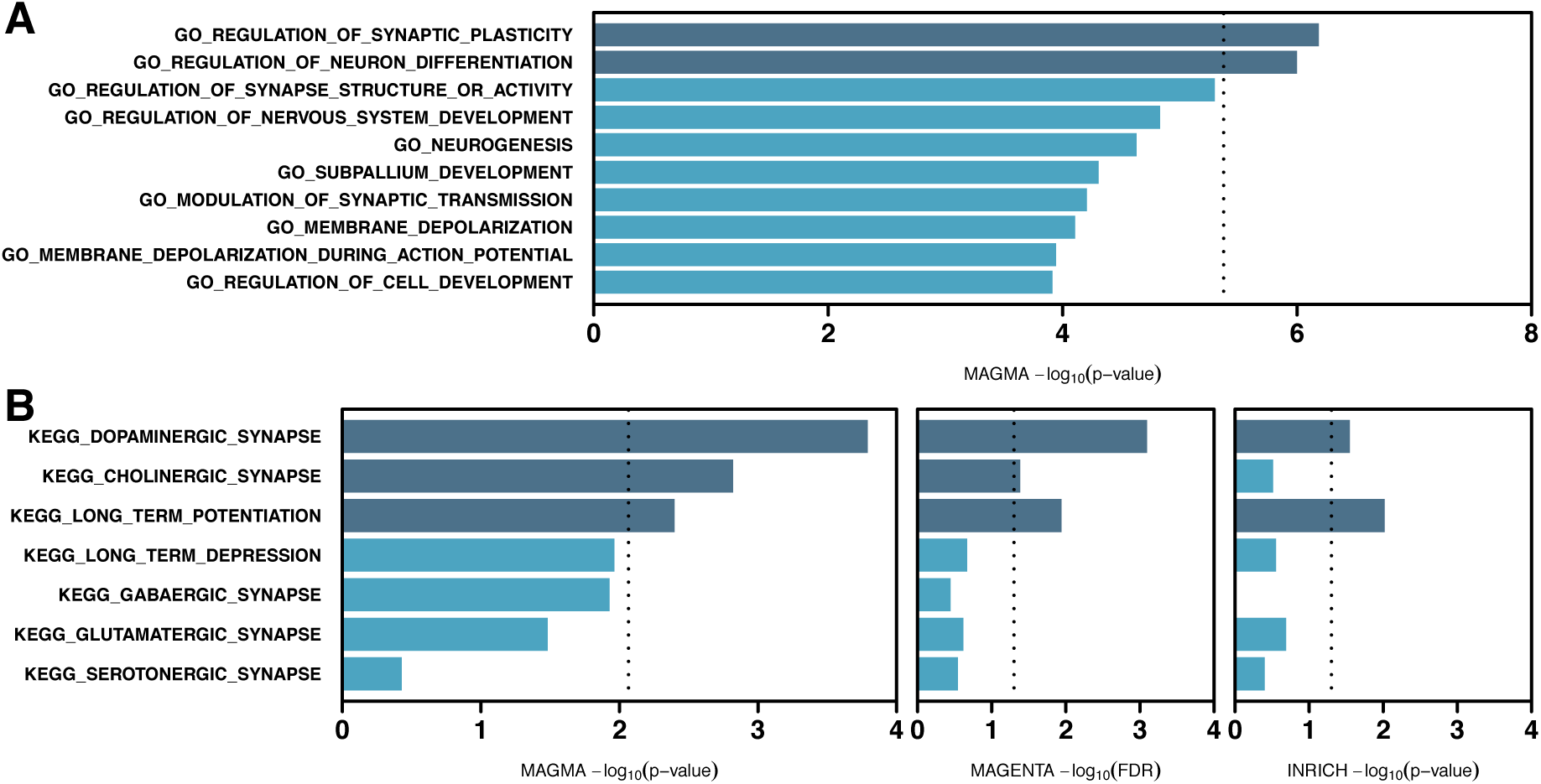
Results of enrichment analysis on MSigDB biological processes and KEGG pathways. Reported p-values or FDR are –log_10_ converted. Significance thresholds are different per analysis depending on the used method. Significant enrichment is indicated with dark blue. **(A)** Top ten enriched MSigDB gene sets (Significance threshold p = 4.23×10^-6^). **(B)** Enrichment in KEGG synaptic signaling pathways tested using MAGMA (left panel, significance threshold p = 8.6×10^-3^), MAGENTA (middle panel, significance threshold FDR = 0.05) and INRICH (right panel, significance threshold p = 0.05).

To gain a more nuanced understanding of molecular synaptic pathways enriched for schizophrenia-associated variants, we tested enrichment of SNPs in KEGG pathways representing synaptic signaling. We found significant enrichment in pathways representing the dopaminergic synapse (p = 1.62×10^-4^), cholinergic synapse (p = 1.51×10^-3^) and long-term potentiation (p = 4.00×10^-3^) (Figure 2B, Supplementary Table S2). For each significant pathway, we mapped top genes (from the MAGMA gene analysis step, see Supplementary Table S3) on components within these KEGG pathways (Figure 3A, Supplementary Figure S5). SNP enrichment was mostly restricted to trans-membrane and postsynaptic components in the cholinergic and dopaminergic synapses. The long-term potentiation pathway only included post-synaptic components. We detected strong enrichment in signaling through extracellular signal-regulated kinase (ERK) and cAMP response element-binding protein (CREB), phospholipase C (PLC) and the inositol trisphosphate receptor (IP_3_R), and protein kinase B (PKB/Akt). These cascades converge on mechanisms involved in synaptic growth regulation and synaptic plasticity. Furthermore, voltage-gated calcium channels, glutamatergic NMDA and AMPA receptors, the dopamine D2 receptor (DRD2) and the muscarinic acetylcholine receptor M_4_ were highly enriched. A proportion of genes was shared between the tested KEGG pathways and enriched GO gene sets (Supplementary Figure S1B). Conditioning the analysis of the two significant GO gene sets on shared genes revealed a slight effect of these genes on the enrichment signal (Supplementary Figure S4A). When we conditioned the analysis of each KEGG pathway on shared genes with other tested KEGG pathways, all enrichment p-values increased to non-significance (p > 8.6×10^-3^), indicating a considerable contribution of shared genes to the enrichment signal (Supplementary Figure S4B). Shared genes between KEGG pathways that were tested as a single gene set were however not enriched after multiple testing correction in a combined analysis with sets of unique genes per pathway (p = 9.4×10^-3^) (Supplementary Figure S6). Protein products of 22 unique top genes from enriched KEGG pathways showed more direct interactions than expected by chance (p = 1.4×10^-2^) and more indirect connections to other proteins (p = 4.0×10^-3^), but not direct connections to other proteins (p = 0.81) (Figure 3B, C). Gene expression analysis revealed no large differences in transcript abundance for top genes among brain tissues, although relative expression seems higher in cerebellum and cerebellar hemisphere compared to other brain tissues (Supplementary Figure S7), and between brain tissues and other tissues (Supplementary Figure S8). When we applied median gene expression levels in the brain as a covariate in the MAGMA competitive gene set analysis of GO gene sets and KEGG pathways, we observed a slight attenuation of the enrichment signal although all enriched GO gene sets and KEGG pathways remained significant (Supplementary Figure S9).

**Figure 3.**
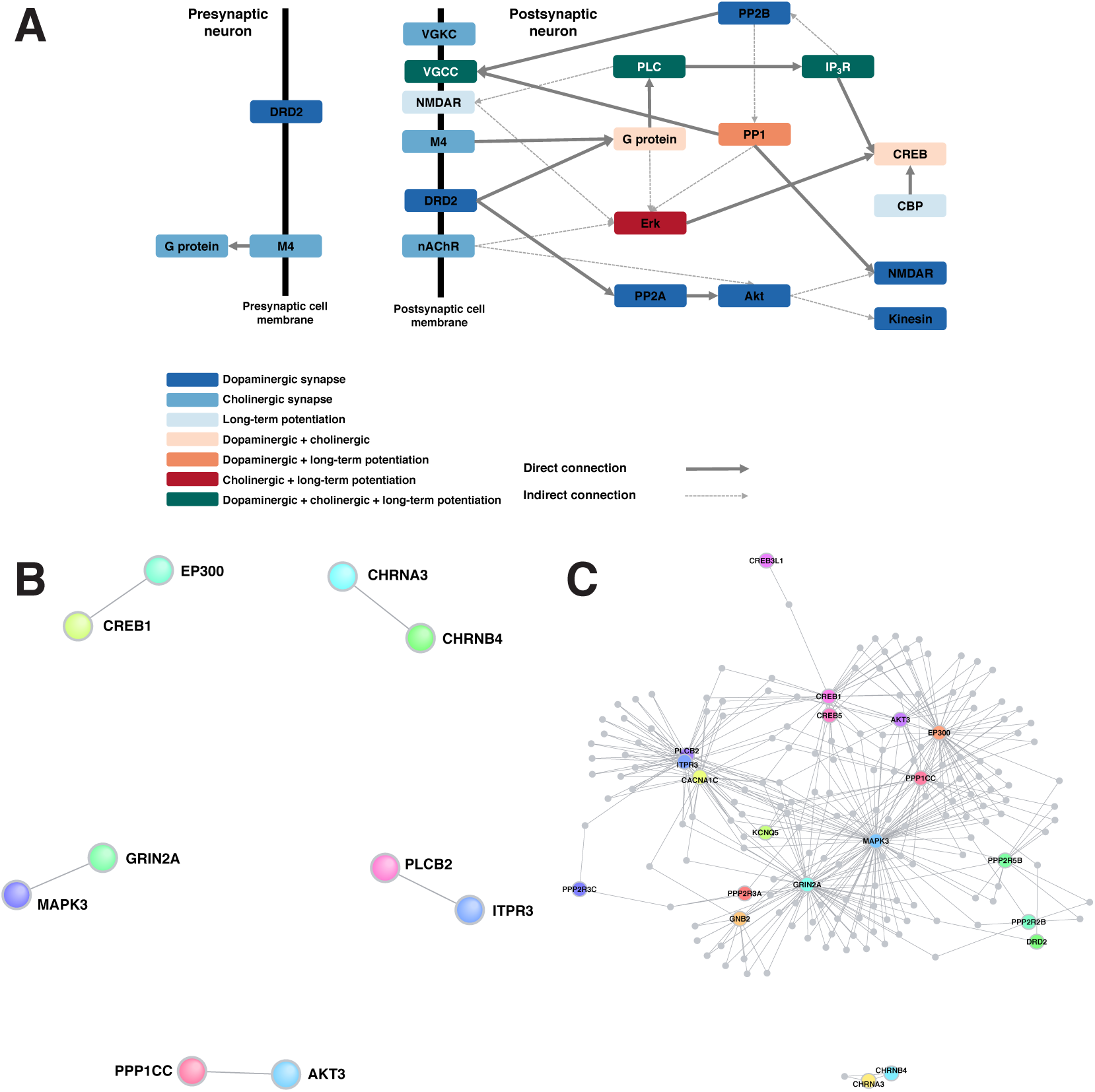
Enrichment and protein-protein interactions in KEGG synaptic signaling pathways. **(A)** Significant pathway components (gene p-value < 1.5×10^-4^) for dopaminergic synapse, cholinergic synapse and long-term potentiation. Colors indicate specificity to one or more pathways. Arrows indicate connections between signaling components (solid, direct interaction; dashed, indirect interaction), based on KEGG.(Kanehisa and Goto, 2000) **(B)** Direct (interaction p = 1.4×10^-2^) and **(C)** indirect (interaction p = 4.0×10^-3^) protein-protein interaction networks generated with DAPPLE (Rossin et al., 2011) for 22 top genes from the dopaminergic synapse, long-term potentiation and cholinergic synapse gene sets.

Using MAGENTA and INRICH with the same data and gene sets, we confirmed significant enrichment of the dopaminergic synapse (MAGENTA corrected FDR = 8.00×10^-4^, INRICH corrected p-value = 2.82×10^-2^) and long-term potentiation (MAGENTA corrected FDR = 1.14×10^-2^, INRICH corrected p-value = 9.6×10^-3^). Enrichment for the cholinergic synapse was only confirmed in MAGENTA (corrected FDR = 4.12×10^-2^) (Figure 2B). When we applied our analysis pipeline to the data of the 2018 schizophrenia GWAS meta-analysis (Pardiñas et al., 2018), we confirmed enrichment of neurodevelopmental processes and the dopaminergic synapse, while results again pointed towards enrichment of the cholinergic synapse. In addition, we also found enrichment of the GABAergic synapse in the MAGMA analysis. Only enrichment in the dopaminergic synapse was confirmed by all three gene set analysis tools (Supplementary Figure S10).

## 4. Discussion

By implementing complementary gene set enrichment analysis tools (MAGMA, MAGENTA and INRICH), annotations from biological databases (MSigDB/GO and KEGG), protein-protein interaction data, and tissue-specific gene expression into a comprehensive analysis, we aimed to elucidate biological processes underlying schizophrenia. Thus, we first detected enrichment of schizophrenia-associated SNPs in synaptic plasticity and neuronal differentiation processes, which is in line with biological hypotheses introduced previously (Hertzberg et al., 2015; Lips et al., 2012; Network and Pathway Analysis Subgroup of Psychiatric Genomics Consortium, 2015). We did however not replicate significant enrichment of immune and histone pathways (Supplementary Figure S11), which have also been reported as highly associated to psychiatric disease (Network and Pathway Analysis Subgroup of Psychiatric Genomics Consortium, 2015). Possible reasons for this lack of replication are the larger sample size and the exclusion of other phenotypes than schizophrenia in the current study. We followed our first findings up in a targeted analysis on pathways representing synaptic signaling in all major neurotransmitter systems and demonstrated enrichment of schizophrenia SNPs in the dopaminergic synapse, long-term potentiation through the glutamatergic system and the cholinergic synapse, although the latter could not be confirmed using INRICH. We then showed that protein products of significant schizophrenia-associated genes in our enriched KEGG pathways show on average a higher rate of interactions with other proteins than expected by chance. Finally, we highlight that expression patterns of top genes from the enriched KEGG pathways do not substantially differ between brain regions and that enrichment levels are not substantially attenuated by including brain gene expression as a covariate in our MAGMA analysis.

Dysfunctional synaptic transmission impacts synaptic plasticity and brain development, mediated through long-term potentiation (LTP) and long-term depression (LTD) (Bernardinelli et al., 2014). Although all five major neurotransmitter systems (dopamine, gamma-aminobutyric acid, glutamate, serotonin, and acetylcholine) have been implicated in schizophrenia, the extent to which each of them is involved had remained elusive (Kahn et al., 2015; Pocklington et al., 2014). Our results strongly support the involvement of the dopaminergic system, which has been extensively examined in schizophrenia. Previous studies have reported increased dopamine synthesis and release, and increased dopamine receptor expression in schizophrenia (Kaalund et al., 2014; Kahn et al., 2015). *DRD2* genetic variants are also implicated in schizophrenia and several of its intermediate phenotypes (Luykx et al., 2017; Vink et al., 2016). We here confirm an accumulation of *DRD2* genetic variants in schizophrenia and in signaling cascades downstream of this receptor. Enrichment of the glutamate-induced LTP pathway was another finding that could be verified using all enrichment analysis tools. Mediation of LTP is, however, not limited to the glutamatergic system as post-synaptic signaling molecules such as the above mentioned CREB, IP_3_R and PKB mediate synaptic plasticity in other neurotransmitter systems (e.g. the dopaminergic system). Multiple lines of evidence link LTP to cognitive deficits in schizophrenia (Salavati et al., 2015). Cholinergic transmission may also be relevant to symptomatology of schizophrenia, especially in light of the high rates of nicotine abuse and a range of cognitive symptoms (Carruthers et al., 2015; Parikh et al., 2016) which have led some to postulate that schizophrenia is primarily a cognitive illness (Kahn and Keefe, 2013). The implication of acetylcholine in schizophrenia is further supported by a landmark study investigating chromatin interactions between enhancer regions containing schizophrenia-associated loci and promoter regions of target genes (Won et al., 2016). In a broader perspective, our findings are in line with rare variant studies in schizophrenia that report mutations in genes coding for postsynaptic signaling components (CNV and Schizophrenia Working Groups of the Psychiatric Genomics Consortium, 2017; Fromer et al., 2014; Kirov et al., 2012; Purcell et al., 2014).

Our detailed analysis of downstream signaling cascades in all major neurotransmitter system gene sets revealed several of these cascades to be highly enriched for schizophrenia-associated variants: the phospholipase pathway, CREB signaling and the PKB/Akt signaling cascade. All of these cascades may be linked to schizophrenia by numerous lines of neurobiological evidence, as outlined below. First, the phospholipase pathway (particularly PLC) controls neuronal activity and thereby maintains synaptic functioning and development. Gene deletions in *PLC* are associated with schizophrenia and altered expression of *Plc* and schizophrenia-like behavior have been reported in *Plc* knock-out mice (Koh, 2013; Vasco et al., 2012). Second, signaling through CREB modulates synaptic plasticity. A recent study focusing on the cyclic adenosine monophosphate (cAMP)/PKA/CREB pathway shows a significant association of a SNP in this system with schizophrenia (Forero et al., 2016). Additionally, ERK is part of the CREB signaling cascade and has been found to be enriched in our analyses. Impairment of signaling through ERK is hypothesized to constitute a disease mechanism in schizophrenia (Kyosseva, 2004; Yuan et al., 2010). Third, we found a significant enrichment of schizophrenia SNPs in postsynaptic protein kinase B (PKB or Akt). *AKT1* messenger RNA levels are higher in blood of schizophrenia patients compared to healthy controls and interactions between genetic variation in *AKT1* and cannabis use are associated with schizophrenia, possibly mediated through AKT signaling downstream of DRD2 (Liu et al., 2016; van Winkel, 2011). Interestingly, phosphorylation of glycogen synthase kinase 3 beta (Gsk3β) by the antipsychotic aripiprazole is mediated by Akt (Pan et al., 2015). Finally, we detected an accumulation of SNPs in protein phosphatase 1 (PP1) and protein phosphatase 2A (PP2A). PP2A is one of the mediators of sensorimotor gating, an intermediate phenotype for schizophrenia (Kapfhamer et al., 2010).

Conditional analysis correcting for overlapping genes between gene sets and pathways indicated a high sharing of genes contributing to enrichment between the two GO gene sets. That these GO gene sets are overlapping and were the only two being significantly enriched, implies a highly specific signal found through this analysis in MAGMA. Furthermore, these gene sets connect well to the analyzed KEGG pathways. Between KEGG pathways, conditioning on shared genes resulted in enrichment significance levels dropping to non-significance, supporting the hypothesis that signaling mechanisms shared between the implicated pathways are likely to be a strong underlying factor in schizophrenia in addition to neurotransmitter-specific pathways. Additionally, the *in silico* finding of higher interaction rates of proteins coded by SNP-enriched genes in the dopaminergic, LTP and cholinergic pathways hint at an important biological role of these genes in various biological processes, including synaptic signaling. Although this interaction may be expected since these proteins all operate in the same cellular compartment (the synapse). Expression profiles of top genes from enriched KEGG pathways among brain tissues did not show major differences, similarly to brain tissues versus other tissues (Supplementary Figure S8). This could support the widespread role of these genes in biological processes. Seemingly higher relative transcript abundance of these genes in cerebellum and cerebellar hemisphere might be explained by the fact that these genes were selected from neuronal gene sets and that neuron density is higher in these tissues compared to the rest of the brain (Azevedo et al., 2009). The relative lack of attenuation of enrichment in GO gene sets and KEGG pathways when conditioned on brain gene expression might not have been expected given that these pathways represent processes and systems in the brain. However, genes with low expression levels can still have a biologically relevant function. Furthermore, gene expression in the GTEx database has been quantified in miscellaneous samples, in whom gene expression levels could differ from a sample limited to schizophrenia subjects. Finally, these gene expression levels were derived from whole brain tissue containing different cell types, whereas our tested pathways only represented a specific cellular compartment.

Several limitations should be considered when interpreting our results. First, as our analyses are dependent on the power of GWASs, we cannot rule out the possibility that increased sample sizes in future studies may flag other synaptic systems also hypothesized to be associated with schizophrenia, such as the glutamatergic system (Yin et al., 2012), as further illustrated by results pointing towards enrichment in the GABAergic synapse in novel GWAS (Pardiñas et al., 2018) (Supplementary Figure S10). Second, we can only test for enrichment in gene sets and pathways that are annotated based on the knowledge currently available. Third, only protein-coding regions of the genome and up- and downstream regions in close proximity to genes were considered in our analyses. Non-coding stretches of the genome account for a major part of disease heritability and transcription regulation (Won et al., 2016). Advances in annotation of the non-coding parts of the genome will likely allow for future integration of these regions in currently available gene set enrichment analysis tools and novel functional genomics tools. These may be integrated with expression quantitative trait locus (eQTL) data and genomic interactions, which potentially prime novel molecular mechanisms involved in the pathways underlying schizophrenia. Finally, MAGMA, MAGENTA and INRICH slightly differ in their approach of testing a generally similar null hypothesis - that genes in a given gene set are not more strongly associated with schizophrenia than genes outside such a gene set. Despite their disparaging analytical approaches but perhaps owing to these similar null hypotheses, we found highly similar results using these tools as would be expected.

In conclusion, using complementary enrichment analysis approaches, we highlight downstream signaling cascades as the most likely part of the dopaminergic, cholinergic and LTP systems to have a role in schizophrenia. Our results open avenues for further research aimed at elucidating signaling pathways in schizophrenia, e.g. through more comprehensive integration with genotype, expression and protein-interaction data. Finally, our findings may aid the discovery of novel drug targets by prioritizing neurotransmitter systems and their signaling molecules, e.g. through translational experiments or drug-interaction studies, to hopefully reduce the burden imposed on quality of life in patients suffering from this disabling disorder.

## Abbreviations

DRD2: dopamine receptor D2
M4: muscarinic acetylcholine receptor M4
G protein: guanine nucleotide-binding protein
VGKC: voltage-gated potassium channel
VGCC: voltage-gated calcium channel
NMDAR: N-methyl-D-aspartate receptor
nAChR: nicotinic acetylcholine receptor
PP2A: protein phosphatase 2A
Akt: protein kinase B
Erk: Extracelluar signal-regulated kinase
CREB: cAMP response element binding protein
CBP: CREB-binding protein
PP1: protein-phosphatase 1
PLC: phospholipase C
IP_3_R: inositol trisphosphate receptor
PP2B: protein-phosphatase 2B

## References

Azevedo, F.A.C., Carvalho, L.R.B., Grinberg, L.T., Farfel, J.M., Ferretti, R.E.L., Leite, R.E.P., Jacob Filho, W., Lent, R., Herculano-Houzel, S., 2009. Equal numbers of neuronal and nonneuronal cells make the human brain an isometrically scaled-up primate brain. J. Comp. Neurol. 513, 532–541. doi:10.1002/cne.21974

Bernardinelli, Y., Nikonenko, I., Muller, D., 2014. Structural plasticity: mechanisms and contribution to developmental psychiatric disorders. Frontiers in Neuroanatomy 8. doi:10.3389/fnana.2014.00123

Carruthers, S.P., Gurvich, C.T., Rossell, S.L., 2015. The muscarinic system, cognition and schizophrenia. Neurosci Biobehav Rev 55, 393–402. doi:10.1016/j.neubiorev.2015.05.011

Chang, C.C., Chow, C.C., Tellier, L.C., Vattikuti, S., Purcell, S.M., Lee, J.J., 2015. Second-generation PLINK: rising to the challenge of larger and richer datasets. GigaScience 4, 7. doi:10.1186/s13742-015-0047-8

CNV and Schizophrenia Working Groups of the Psychiatric Genomics Consortium, 2017. Contribution of copy number variants to schizophrenia from a genome-wide study of 41,321 subjects. Nature genetics 49, 27–35. doi:10.1038/ng.3725

de Leeuw, C.A., Mooij, J.M., Heskes, T., Posthuma, D., 2015. MAGMA: Generalized Gene-Set Analysis of GWAS Data. PLoS Comput Biol 11, e1004219. doi:10.1371/journal.pcbi.1004219

de Leeuw, C.A., Neale, B.M., Heskes, T., Posthuma, D., 2016. The statistical properties of gene-set analysis. Nat. Rev. Genet. 17, 353–364. doi:10.1038/nrg.2016.29

Duncan, L.E., Holmans, P.A., Lee, P.H., O’Dushlaine, C.T., Kirby, A.W., Smoller, J.W., Öngür, D., Cohen, B.M., 2014. Pathway analyses implicate glial cells in schizophrenia. PloS one 9, e89441. doi:10.1371/journal.pone.0089441

Forero, D.A., Herteleer, L., De Zutter, S., Norrback, K.F., Nilsson, L.G., Adolfsson, R., Callaerts, P., Del-Favero, J., 2016. A network of synaptic genes associated with schizophrenia and bipolar disorder. Schizophrenia research 172, 68–74. doi:10.1016/j.schres.2016.02.012

Fromer, M., Pocklington, A.J., Kavanagh, D.H., Williams, H.J., Dwyer, S., Gormley, P., Georgieva, L., Rees, E., Palta, P., Ruderfer, D.M., Carrera, N., Humphreys, I., Johnson, J.S., Roussos, P., Barker, D.D., Banks, E., Milanova, V., Grant, S.G., Hannon, E., Rose, S.A., Chambert, K., Mahajan, M., Scolnick, E.M., Moran, J.L., Kirov, G., Palotie, A., McCarroll, S.A., Holmans, P., Sklar, P., Owen, M.J., Purcell, S.M., O’Donovan, M.C., 2014. De novo mutations in schizophrenia implicate synaptic networks. Nature 506, 179–184. doi:10.1038/nature12929

Gaspar, H.A., Breen, G., 2017. Drug enrichment and discovery from schizophrenia genome-wide association results: an analysis and visualisation approach. Scientific reports 7, 12460. doi:10.1038/s41598-017-12325-3

Gene Ontology Consortium, 2015. Gene Ontology Consortium: going forward. Nucleic acids research 43, D1049–56. doi:10.1093/nar/gku1179

Hertzberg, L., Katsel, P., Roussos, P., Haroutunian, V., Domany, E., 2015. Integration of gene expression and GWAS results supports involvement of calcium signaling in Schizophrenia. Schizophrenia research 164, 92–99. doi:10.1016/j.schres.2015.02.001

Kaalund, S.S., Newburn, E.N., Ye, T., Tao, R., Li, C., Deep-Soboslay, A., Herman, M.M., Hyde, T.M., Weinberger, D.R., Lipska, B.K., Kleinman, J.E., 2014. Contrasting changes in DRD1 and DRD2 splice variant expression in schizophrenia and affective disorders, and associations with SNPs in postmortem brain. Molecular psychiatry 19, 1258–1266. doi:10.1038/mp.2013.165

Kahn, R.S., Keefe, R.S., 2013. Schizophrenia is a cognitive illness: time for a change in focus. JAMA psychiatry 70, 1107–1112. doi:10.1001/jamapsychiatry.2013.155

Kahn, R.S., Sommer, I.E., Murray, R.M., Meyer-Lindenberg, A., Weinberger, D.R., Cannon, T.D., O’Donovan, M., Correll, C.U., Kane, J.M., van Os, J., Insel, T.R., 2015. Schizophrenia. Nature Reviews Disease Primers 15067. doi:10.1038/nrdp.2015.67

Kanehisa, M., Goto, S., 2000. KEGG: kyoto encyclopedia of genes and genomes. Nucleic acids research 28, 27–30.

Kapfhamer, D., Berger, K.H., Hopf, F.W., Seif, T., Kharazia, V., Bonci, A., Heberlein, U., 2010. Protein Phosphatase 2a and glycogen synthase kinase 3 signaling modulate prepulse inhibition of the acoustic startle response by altering cortical M-Type potassium channel activity. The Journal of neuroscience: the official journal of the Society for Neuroscience 30, 8830–8840. doi:10.1523/jneurosci.1292-10.2010

Kirov, G., Pocklington, A.J., Holmans, P., Ivanov, D., Ikeda, M., Ruderfer, D., Moran, J., Chambert, K., Toncheva, D., Georgieva, L., Grozeva, D., Fjodorova, M., Wollerton, R., Rees, E., Nikolov, I., van de Lagemaat, L.N., Bayés, A., Fernandez, E., Olason, P.I., Böttcher, Y., Komiyama, N.H., Collins, M.O., Choudhary, J., Stefansson, K., Stefansson, H., Grant, S.G.N., Purcell, S., Sklar, P., O’Donovan, M.C., Owen, M.J., 2012. De novo CNV analysis implicates specific abnormalities of postsynaptic signalling complexes in the pathogenesis of schizophrenia. Mol. Psychiatry 17, 142–153. doi:10.1038/mp.2011.154

Koh, H.Y., 2013. Phospholipase C-beta1 and schizophrenia-related behaviors. Advances in biological regulation 53, 242–248. doi:10.1016/j.jbior.2013.08.002

Kyosseva, S.V., 2004. The role of the extracellular signal-regulated kinase pathway in cerebellar abnormalities in schizophrenia. Cerebellum 3, 94–99. doi:10.1080/14734220410029164

Lee, P.H., O’Dushlaine, C., Thomas, B., Purcell, S.M., 2012. INRICH: interval-based enrichment analysis for genome-wide association studies. Bioinformatics (Oxford, England) 28, 1797–1799. doi:10.1093/bioinformatics/bts191

Liberzon, A., Subramanian, A., Pinchback, R., Thorvaldsdottir, H., Tamayo, P., Mesirov, J.P., 2011. Molecular signatures database (MSigDB) 3.0. Bioinformatics (Oxford, England) 27, 1739–1740. doi:10.1093/bioinformatics/btr260

Lichtenstein, P., Björk, C., Hultman, C.M., Scolnick, E., Sklar, P., Sullivan, P.F., 2006. Recurrence risks for schizophrenia in a Swedish national cohort. Psychological medicine 36, 1417–1425. doi:10.1017/S0033291706008385

Lips, E.S., Cornelisse, L.N., Toonen, R.F., Min, J.L., Hultman, C.M., Holmans, P.A., O’Donovan, M.C., Purcell, S.M., Smit, A.B., Verhage, M., Sullivan, P.F., Visscher, P.M., Posthuma, D., 2012. Functional gene group analysis identifies synaptic gene groups as risk factor for schizophrenia. Molecular psychiatry 17, 996–1006. doi:10.1038/mp.2011.117

Liu, L., Luo, Y., Zhang, G., Jin, C., Zhou, Z., Cheng, Z., Yuan, G., 2016. The mRNA expression of DRD2, PI3KCB, and AKT1 in the blood of acute schizophrenia patients. Psychiatry research 243, 397–402. doi:10.1016/j.psychres.2016.07.010

Lonsdale, J., Thomas, J., Salvatore, M., Phillips, R., Lo, E., Shad, S., Hasz, R., Walters, G., Garcia, F., Young, N., Foster, B., Moser, M., Karasik, E., Gillard, B., Ramsey, K., Sullivan, S., Bridge, J., Magazine, H., Syron, J., Fleming, J., Siminoff, L., Traino, H., Mosavel, M., Barker, L., Jewell, S., Rohrer, D., Maxim, D., Filkins, D., Harbach, P., Cortadillo, E., Berghuis, B., Turner, L., Hudson, E., Feenstra, K., Sobin, L., Robb, J., Branton, P., Korzeniewski, G., Shive, C., Tabor, D., Qi, L., Groch, K., Nampally, S., Buia, S., Zimmerman, A., Smith, A., Burges, R., Robinson, K., Valentino, K., Bradbury, D., Cosentino, M., Diaz-Mayoral, N., Kennedy, M., Engel, T., Williams, P., Erickson, K., Ardlie, K., Winckler, W., Getz, G., DeLuca, D., MacArthur, D., Kellis, M., Thomson, A., Young, T., Gelfand, E., Donovan, M., Meng, Y., Grant, G., Mash, D., Marcus, Y., Basile, M., Liu, J., Zhu, J., Tu, Z., Cox, N.J., Nicolae, D.L., Gamazon, E.R., Im, H.K., Konkashbaev, A., Pritchard, J., Stevens, M., Flutre, T., Wen, X., Dermitzakis, E.T., Lappalainen, T., Guigo, R., Monlong, J., Sammeth, M., Koller, D., Battle, A., Mostafavi, S., McCarthy, M., Rivas, M., Maller, J., Rusyn, I., Nobel, A., Wright, F., Shabalin, A., Feolo, M., Sharopova, N., Sturcke, A., Paschal, J., Anderson, J.M., Wilder, E.L., Derr, L.K., Green, E.D., Struewing, J.P., Temple, G., Volpi, S., Boyer, J.T., Thomson, E.J., Guyer, M.S., Ng, C., Abdallah, A., Colantuoni, D., Insel, T.R., Koester, S.E., Little, A.R., Bender, P.K., Lehner, T., Yao, Y., Compton, C.C., Vaught, J.B., Sawyer, S., Lockhart, N.C., Demchok, J., Moore, H.F., 2013. The Genotype-Tissue Expression (GTEx) project. Nature genetics 45, 580–585. doi:10.1038/ng.2653

Luo, W., Brouwer, C., 2013. Pathview: an R/Bioconductor package for pathway-based data integration and visualization. Bioinformatics 29, 1830–1831. doi:10.1093/bioinformatics/btt285

Luykx, J.J., Broersen, J.L., de Leeuw, M., 2017. The DRD2 rs1076560 polymorphism and schizophrenia-related intermediate phenotypes: A systematic review and meta-analysis. Neurosci Biobehav Rev 74, Part A, 214–224. doi:10.1016/j.neubiorev.2017.01.006

Network and Pathway Analysis Subgroup of Psychiatric Genomics Consortium, 2015. Psychiatric genome-wide association study analyses implicate neuronal, immune and histone pathways. Nature neuroscience 18, 199–209. doi:10.1038/nn.3922

Pan, B., Chen, J., Lian, J., Huang, X.-F., Deng, C., 2015. Unique Effects of Acute Aripiprazole Treatment on the Dopamine D2 Receptor Downstream cAMP-PKA and Akt-GSK3ß Signalling Pathways in Rats. PloS one 10, e0132722. doi:10.1371/journal.pone.0132722

Pardiñas, A.F., Holmans, P., Pocklington, A.J., Escott-Price, V., Ripke, S., Carrera, N., Legge, S.E., Bishop, S., Cameron, D., Hamshere, M.L., Han, J., Hubbard, L., Lynham, A., Mantripragada, K., Rees, E., MacCabe, J.H., McCarroll, S.A., Baune, B.T., Breen, G., Byrne, E.M., Dannlowski, U., Eley, T.C., Hayward, C., Martin, N.G., McIntosh, A.M., Plomin, R., Porteous, D.J., Wray, N.R., Caballero, A., Geschwind, D.H., Huckins, L.M., Ruderfer, D.M., Santiago, E., Sklar, P., Stahl, E.A., Won, H., Agerbo, E., Als, T.D., Andreassen, O.A., Bækvad-Hansen, M., Mortensen, P.B., Pedersen, C.B., Børglum, A.D., Bybjerg-Grauholm, J., Djurovic, S., Durmishi, N., Pedersen, M.G., Golimbet, V., Grove, J., Hougaard, D.M., Mattheisen, M., Molden, E., Mors, O., Nordentoft, M., Pejovic-Milovancevic, M., Sigurdsson, E., Silagadze, T., Hansen, C.S., Stefansson, K., Stefansson, H., Steinberg, S., Tosato, S., Werge, T., GERAD1 Consortium:, Collier, D.A., Rujescu, D., Kirov, G., Owen, M.J., O’Donovan, M.C., Walters, J.T.R., CRESTAR Consortium:, 2018. Common schizophrenia alleles are enriched in mutation-intolerant genes and in regions under strong background selection. Nat. Genet. 199, 441. doi:10.1038/s41588-018-0059-2

Parikh, V., Kutlu, M.G., Gould, T.J., 2016. nAChR dysfunction as a common substrate for schizophrenia and comorbid nicotine addiction: Current trends and perspectives. Schizophrenia research 171, 1–15. doi:10.1016/j.schres.2016.01.020

Pocklington, A.J., O’Donovan, M., Owen, M.J., 2014. The synapse in schizophrenia. Eur J Neurosci 39, 1059–1067. doi:10.1111/ejn.12489

Purcell, S.M., Moran, J.L., Fromer, M., Ruderfer, D., Solovieff, N., Roussos, P., O’Dushlaine, C., Chambert, K., Bergen, S.E., Kähler, A., Duncan, L., Stahl, E., Genovese, G., Fernández, E., Collins, M.O., Komiyama, N.H., Choudhary, J.S., Magnusson, P.K.E., Banks, E., Shakir, K., Garimella, K., Fennell, T., DePristo, M., Grant, S.G.N., Haggarty, S.J., Gabriel, S., Scolnick, E.M., Lander, E.S., Hultman, C.M., Sullivan, P.F., McCarroll, S.A., Sklar, P., 2014. A polygenic burden of rare disruptive mutations in schizophrenia. Nature 506, 185–190. doi:10.1038/nature12975

Rossin, E.J., Lage, K., Raychaudhuri, S., Xavier, R.J., Tatar, D., Benita, Y., International Inflammatory Bowel Disease Genetics Constortium, Cotsapas, C., Daly, M.J., 2011. Proteins encoded in genomic regions associated with immune-mediated disease physically interact and suggest underlying biology. PLoS Genet. 7, e1001273. doi:10.1371/journal.pgen.1001273

Salavati, B., Rajji, T.K., Price, R., Sun, Y., Graff-Guerrero, A., Daskalakis, Z.J., 2015. Imaging-based neurochemistry in schizophrenia: a systematic review and implications for dysfunctional long-term potentiation. Schizophrenia bulletin 41, 44–56. doi:10.1093/schbul/sbu132

Schizophrenia Working Group of the Psychiatric Genomics Consortium, 2014. Biological insights from 108 schizophrenia-associated genetic loci. Nature 511, 421–427. doi:10.1038/nature13595

Segrè, A.V., Groop, L., Mootha, V.K., Daly, M.J., Altshuler, D., DIAGRAM Consortium, MAGIC investigators, 2010. Common Inherited Variation in Mitochondrial Genes Is Not Enriched for Associations with Type 2 Diabetes or Related Glycemic Traits. PLoS Genet 6, e1001058. doi:10.1371/journal.pgen.1001058

The 1000 Genomes Project Consortium, C., 2015. A global reference for human genetic variation. Nature 526, 68–74. doi:10.1038/nature15393

van Winkel, R., 2011. Family-based analysis of genetic variation underlying psychosis-inducing effects of cannabis: sibling analysis and proband follow-up. Archives of general psychiatry 68, 148–157. doi:10.1001/archgenpsychiatry.2010.152

Vasco, Lo, V.R., Cardinale, G., Polonia, P., 2012. Deletion of PLCB1 gene in schizophrenia-affected patients. Journal of cellular and molecular medicine 16, 844–851. doi:10.1111/j.1582-4934.2011.01363.x

Veyrieras, J.B., Kudaravalli, S., Kim, S.Y., Dermitzakis, E.T., Gilad, Y., Stephens, M., Pritchard, J.K., 2008. High-resolution mapping of expression-QTLs yields insight into human gene regulation. PLoS genetics 4, e1000214. doi:10.1371/journal.pgen.1000214

Vink, M., de Leeuw, M., Luykx, J.J., van Eijk, K.R., van den Munkhof, H.E., van Buuren, M., Kahn, R.S., 2016. DRD2 Schizophrenia-Risk Allele Is Associated With Impaired Striatal Functioning in Unaffected Siblings of Schizophrenia Patients. Schizophrenia bulletin 42, 843–850. doi:10.1093/schbul/sbv166

Wang, K., Gaitsch, H., Poon, H., Cox, N.J., Rzhetsky, A., 2017. Classification of common human diseases derived from shared genetic and environmental determinants. Nat. Genet. 6, 111. doi:10.1038/ng.3931

Warnes, G.R., Ben Bolker, Bonebakker, L., Gentleman, R., Huber, W., Liaw, A., Lumley, T., Maechler, M., Magnusson, A., Moeller, S., Schwartz, M., Venables, B., n.d. gplots: Various R Programming Tools for Plotting Data. R package version 3.0.1. [WWW Document]. URL https://CRAN.R-project.org/package=gplots (accessed 11.6.17).

Won, H., la Torre-Ubieta, de, L., Stein, J.L., Parikshak, N.N., Huang, J., Opland, C.K., Gandal, M.J., Sutton, G.J., Hormozdiari, F., Lu, D., Lee, C., Eskin, E., Voineagu, I., Ernst, J., Geschwind, D.H., 2016. Chromosome conformation elucidates regulatory relationships in developing human brain. Nature 538, 523–527. doi:10.1038/nature19847

Yin, D.M., Chen, Y.J., Sathyamurthy, A., Xiong, W.C., Mei, L., 2012. Synaptic dysfunction in schizophrenia. Advances in experimental medicine and biology 970, 493–516. doi:10.1007/978-3-7091-0932-8_22

Yuan, P., Zhou, R., Wang, Y., Li, X., Li, J., Chen, G., Guitart, X., Manji, H.K., 2010. Altered levels of extracellular signal-regulated kinase signaling proteins in postmortem frontal cortex of individuals with mood disorders and schizophrenia. J Affect Disord 124, 164–169. doi:10.1016/j.jad.2009.10.017

